# Widespread cytoplasmic polyadenylation programs asymmetry in the germline and early embryo

**DOI:** 10.1101/428540

**Authors:** Peter R. Boag, Paul F. Harrison, Adele A. Barugahare, Andrew D. Pattison, Angavai Swaminathan, Greta Raymant, Stephanie Monk, Kirill Tsyganov, Eva Heinz, Gregory M. Davis, David R. Powell, Traude H. Beilharz

## Abstract

**BACKGROUND:** The program of embryonic development is launched by selective activation of a silent maternal transcriptome. In *Caenorhabditis elegans*, nuclei of the adult germline are responsible for the synthesis of at least two distinct mRNA populations; those required for housekeeping functions, and those that program the oocyte-to-embryo transition. We mapped this separation by changes to the length-distribution of poly(A)-tails that depend on GLD-2 mediated cytoplasmic polyadenylation and its regulators genome-wide.

**RESULTS:** More than 1000 targets of cytoplasmic polyadenylation were identified by differential polyadenylation. Amongst mRNA with the greatest dependence on GLD-2 were those encoding RNA binding proteins with known roles in spatiotemporal patterning such as *mex-5* and *pos-1*. In General, the 3’ UTR of GLD-2 targets were longer, contained cytosine-patches, and were enriched for non-standard polyadenylation-motifs. To identify the deadenylase that initiated transcript silencing, we depleted the known deadenylases in the *gld-2(0)* mutant background. Only the loss of CCF-1 suppressed the short-tailed phenotype of GLD-2 targets suggesting that in addition to its general role in RNA turnover, this is the major deadenylase for regulatory silencing of maternal mRNA. Analysis of poly(A)-tail length-change in the embryo lacking specific RNA-binding proteins revealed new candidates for asymmetric expression in the first embryonic divisions.

**CONCLUSION:** The concerted action of RNA binding proteins exquisitely regulates GLD-2 activity in space and time. We present our data as interactive web resources for a model where GLD-2 mediated cytoplasmic polyadenylation regulates target mRNA at each stage of worm germline and early embryonic development.

## Background

The road from transcriptome to proteome is not a linear path. Within the crowded dimensions of the cell, there is a balance in time, place and scale for everything. Thus, multiple post-transcriptional regulatory machineries intersect to fine-tune where, when and to what level mRNA is translated. An exemplar of this complexity is the germline syncytium of the nematode *Caenorhabditis elegans*. In this system, transcription and translation of a significant proportion of the transcriptome is separated in both space and time (Fig 1a). The syncytium links a hub of stem cells at one end of the germline, and the developing oocytes and early embryos at the other. Between these distinct cell types are the nuclei that provide mRNA for immediate building and maintenance projects within the syncytium, and a second silent transcriptome that is stored for activation at later timepoints in development. These same nuclei are then transcriptionally silenced for mRNA production during meiosis and remain silenced until the 4-cell embryo stage (1, 2). The steps that drive transcriptional activation of the zygotic genome are thus all post-transcriptionally controlled and are driven by selective retrieval of stored maternal mRNA for translation. An essential driver of this reactivation is the re-extension of the silent, deadenylated mRNA by cytoplasmic polyadenylation.

**Figure 1:**
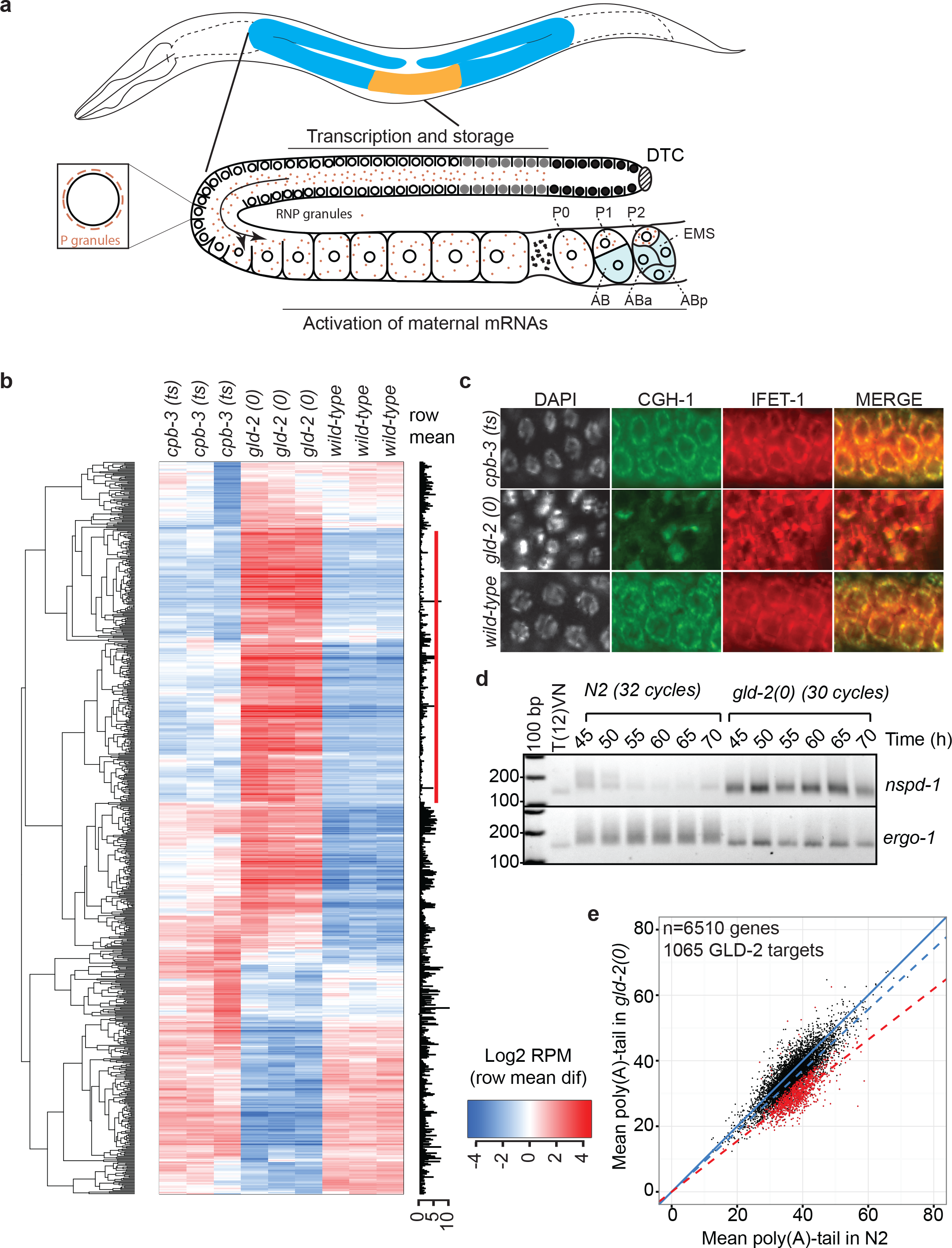
PAT-seq reveals differential expression and polyadenylation-state in the *gld-2(0)* mutant transcriptome. **a)** Schematic representation of the nematode worm *C. elegans* highlighting one arm of the adult hermaphrodite germline. The distal tip cell (DTC) provides the nuclei that populate the shared cytoplasm. The nuclei (open circles) of the upper germline support synthesis of two transcriptomes, one that is utilized to build and maintain the syncytium and one that is captured by P-granules that first abut nuclear pore complexes (see detail), but are later shed into the core of the syncytium and distribute to the developing oocytes and early embryos. The maternal mRNA stored in these granules are activated by the concerted activity of RNA binding proteins along this trajectory of development. The worm specific nomenclature for the first cell divisions is given for context when tracking activation throughout the manuscript. The blue shading of the embryo indicates a somatic cell fate whereas the P2 cell is the germline precursor. **b)** Gene expression changes in *gld-2(0)* mutants. Clustering of raw PAT-seq read-count data for 3 biological replicates of each *cpb-3*, *gld-2* and wild-type worms. Each gene (horizontal line) is colored red for up regulated, or blue down regulated relative to row-mean. Black bars indicate the relative abundance (log_2_ reads per million). The red vertical bar indicates a cluster of transcripts associated with the male germline that are aberrantly retained after switch to the oogenic germline in the adult hermaphrodite worm. **c)** The germline of *gld-2(0)* mutants is disordered. Surface views of the adult germline syncytium stained either with DAPI to localize Nuclear DNA, or immunofluorescence with two markers of perinuclear germ granules CGH-1 (green) and IFET-1 (red). **d)** Retained spermatogenic mRNA in the *gld-2(0)* mutant stabilized with short A-tails. The expression and polyadenylation of *nspd-1*(male) and *ergo-1* (male and female) were monitored over the indicated time-points of development, from late L4 to adulthood, using the ePAT method for detection of polyadenylation. The T(12)VN lane represents the migration of the A-12 form of the PCR amplicon. Any smear of amplicons migrating more slowly than this represent increasing poly(A)-tail length. Note that *nspd-1* required fewer cycles of amplification for detection in the *gld-2(0)* mutant, consistent with its aberrant retention in the adult *gld-2(0)* transcriptome. **e)** PAT-seq detects changes to poly(A)-tail length distribution. The scatter plot represents the mean sequenced adenylation-state of 6510 genes that pass filtering in both the *gld-2(0)* and wild-type transcriptomes. The solid blue line represents tail-length parity, whereas the dashed blue line indicates the actual length-distribution observed. In red, and indicated by the red-dashed line are the 1065 genes that are high-confidence targets of cytoplasmic adenylation (see methods and interactive differential tests (14)).

Cytoplasmic polyadenylation was first identified in studies of early embryonic development in model organisms (reviewed: (3–5). Seminal studies in the sea urchin showed that fertilization resulted in a massive increase in RNA polyadenylation (6). The gene responsible for this post-transcriptional extension of poly(A)-tails was subsequently identified by cloning of the embryonic lethal *gld-2* (for germ line defective) *C. elegans* mutant (7). Since that time, protein homologues have been identified across metazoan lineages (8) and functional roles for cytoplasmic polyadenylation have been extended to span multiple states in development and disease.

It is generally thought that GLD-2 is a constitutively active enzyme, needing only a partner RNA binding protein to anchor it to 3’ UTR ((9) and reviewed: (5)). One such family of RNA binding proteins are the cytoplasmic polyadenylation element (CPE) binding proteins (CPEB). Foremost among these, CPEB1 was shown to control cytoplasmic adenylation and subsequent translation of core cell cycle transcripts (10). Furthermore, the interaction between CPEB1 and its 3’ UTR binding site in target mRNA has been instrumental in building the idea that recognition codes between RNA-binding proteins and 3’ UTR specify translational control (11).

To understand the breadth of control that cytoplasmic adenylation has over post-transcriptional gene expression, we turned to the *C. elegans* model. We reasoned that changes to polyadenylation might identify functionally distinct classes of mRNA required within the germline and in the critical oocyte-to-embryo transition. That is, that we could track initial silencing of mRNA by deadenylation followed by temporal and spatially-distant activation of mRNA by cytoplasmic polyadenylation. Applying the PAT-seq approach to identify global changes to gene expression, 3’ UTR choice and poly(A)-tail length-distribution (12), we searched for mRNA that change in adenylation-state in response to the loss of GLD-2 enzymatic function. This approach identified a large number of transcripts subject to cytoplasmic polyadenylation. Moreover, we show that the short-tailed phenotype of GLD-2 target mRNA is dependent on an initial deadenylation by CCF-1, and occurs on mRNA having a fundamentally different 3’ UTR character than non-target mRNA. We asked if changes to adenylation-state might also be used to identify mRNA responsible for asymmetric division and patterning of the early embryo. Analysis the transcriptome of 1-2 cell embryos highlights many transcripts that become activated at this developmental timepoint. Moreover, reanalysis of previous PAT-seq data from embryos lacking key regulatory RNA-binding proteins (13) revealed multiple transcripts that are subject to asymmetric activation in the early embryo. These data are all available as a resource to the community through a compendium of interactive web-based visualization tools (14).

## Results

### Identifying the targets of GLD-2 mediated cytoplasmic adenylation in *Caenorhabditis elegans*

The *C. elegans* germline provides a genetically tractable model for a number of conserved metazoan pathways of post-transcriptional gene regulation (15, 16). Cytoplasmic polyadenylation has long been considered as an integral feature of this, with GLD-2 activity being essential for fertility (7, 17). However, whether this reflects a dependence of a few key regulators on cytoplasmic polyadenylation, or if it represents a default modification of the germline transcriptome was unknown. To better understand the targeting of cytoplasmic adenylation, we applied the PAT-seq method for identification of changes to 3’-end dynamics (12). By this approach, we expected to identify mRNAs that change in overall expression and in the length-distribution of poly(A)-tails in response to loss of GLD-2 enzymatic activity.

Our first step was to compare the adult transcriptomes of wild-type (N2) worms and those harboring a loss-of-function mutation in GLD-2 (referred to as *gld-2(0)*) (18) and a temperature sensitive allele of CPB-3 (referred to as *cbp-3(ts)*) (19). Our rationale was that mRNA that depend on the enzymatic function of GLD-2 would have shorter poly(A)-tails in the *gld-2(0)* mutant compared to the wild-type transcriptome (7, 17, 20–22). Moreover, we hypothesized that *cpb-3(ts)*, as the closest worm homolog of the CPEB1 recruitment factor for the vertebrate cytoplasmic polyadenylase, might show a similar short-tailed phenotype for at least a subset of germline mRNAs. CPB-3 is one of four CPEB proteins in the worm, and the only one with a reported function in oogenesis (19), the others having described roles in the sex-determination cascades (23). The transcriptomes of one-day-old adult wild-type, *gld-2(0)* and *cbp-3(ts)* worms were analyzed in triplicate.

Global analysis of the transcriptome revealed that many mRNAs were differentially expressed between the three strains (Fig 1b), but overall, that the wild-type and *cpb-3(ts)* transcriptomes were more similar to each other than the *gld-2(0)* mutant. All genome-wide data can be visualized and interactively interrogated at the dedicated companion website (14). Unexpectedly, a number of mRNA encoding sperm-specific proteins were aberrantly retained in *gld-2(0)* adults (Fig 1b, red bar). Although a smaller proportion of such transcripts were also retained in the *cpb-3(ts)* strain, neither this nor any of our subsequent analyses suggested a direct phenocopy of the *gld-2(0)* phenotype by the *cpb-3(ts)* strain. For example, immuno-staining of adult worms to focus on the architecture of the germ cells, showed major disturbances in the *gld-2(0)* mutant, whereas *cpb-3(ts)* mutants looked structurally normal (Fig 1c). We concluded therefore, that in contrast to *gld-2(0)* mutants, the temperature sensitive allele of *cpb-3* did not hold an overt influence over RNA dynamics in the germline when analyzed at the current level of resolution.

### GLD-2 is required for normal metabolism of the spermatogenic transcriptome

The switch from male to female germ-cell development occurs toward the end of the last larval stage (L4) in the worm. This is reflected by a switch from the spermatogenesis program to the oogenic transcriptome. The persistence of spermatogenic transcripts in the one-day-old *gld-2(0)* adult could be due to an aberrant retention of mRNA after the sperm-to-oocyte switch, or to stochastic developmental delay that results in worms at mixed stages of development. Phenotypically, the *gld-2(0)* mutants are female, albeit with a disordered germ-line morphology. However, to rule out any gross delay to development we analyzed the expression and poly(A)-tail length-distribution of a candidate male transcript (*nspd-1*) over a time-course of development from 45-70 hours by the PCR-based ePAT approach to poly(A)-length assessment (22). In wild-type worms, the poly(A)-tail of *nspd-1* presented with a wide length-distribution at 45 hours, which contracted with age and decay during the switch to an oogenic germline at ~55 hours (Fig 1d). Strikingly, in *gld-2(0)* mutants, the transcript was short-tailed and remained abundant throughout the time-course, suggesting that transcript retention was not simply due to gross delay of germ cell development, but rather, a failure of normal metabolism and clearance.

The *Drosophila* homolog of GLD-2, *Wisp*, is necessary for normal sperm development (24). To ask therefore if cytoplasmic adenylation is a common feature of spermatogenic transcripts in *C. elegans*, we compared the average length of the poly(A)-tail for the spermatogenic transcripts (Fig 1b, red bar) in wild-type versus *gld-2(0)* transcriptomes. Of the 177 transcripts in this group, 104 were present with enough reads in the wild-type adult to determine an adenylation-state difference. Whereas the average A-tract sequenced for these transcripts in wild-type was 41, in *gld-2(0)* this reduced to an average of 30 adenosine residues (p=2.05e^−13^); see gene list in additional table 1). To our knowledge, this is the first indication that transcripts of the male germline depend on cytoplasmic polyadenylation in *C. elegans*.

### GLD-2 broadly regulates the oogenic transcriptome

Next, we focused our analysis on the general adenylation-state difference between the adult wild-type and *gld-2(0)* transcriptomes, noting that this is dominated by the oogenic germline. Using a conservative statistical cut-off (FDR < 0.01), we identified 1065 mRNA that vary in poly(A)-tail distribution between wild-type and *gld-2(0)*, with these being overwhelmingly shorter in the mutant than in wild-type ((Fig 1e,); and see interactive differential tests (14)). Amongst transcripts with the most statistically robust shortening were those encoding RNA-binding proteins that have previously been implicated in the orchestration of spatiotemporal expression in *C. elegans* development (13, 25, 26) and (Fig 2a). The large number of transcripts that change in adenylation at a subtler level caused an apparent global shift toward shorter poly(A)-tails (see dashed blue line, Fig 1e). However, focus on such apparent global shifts obscures the fact that multiple functional groups do not change in adenylation-state. For example, transcripts encoding ribosomal proteins did not differ in their adenylation-state between wild-type and *gld-2(0)* worms (Fig 2a). The easy to use web app utilized to generate this figure is available for exploration of further gene ontology terms and/or genes of interest (14).

**Fig 2:**
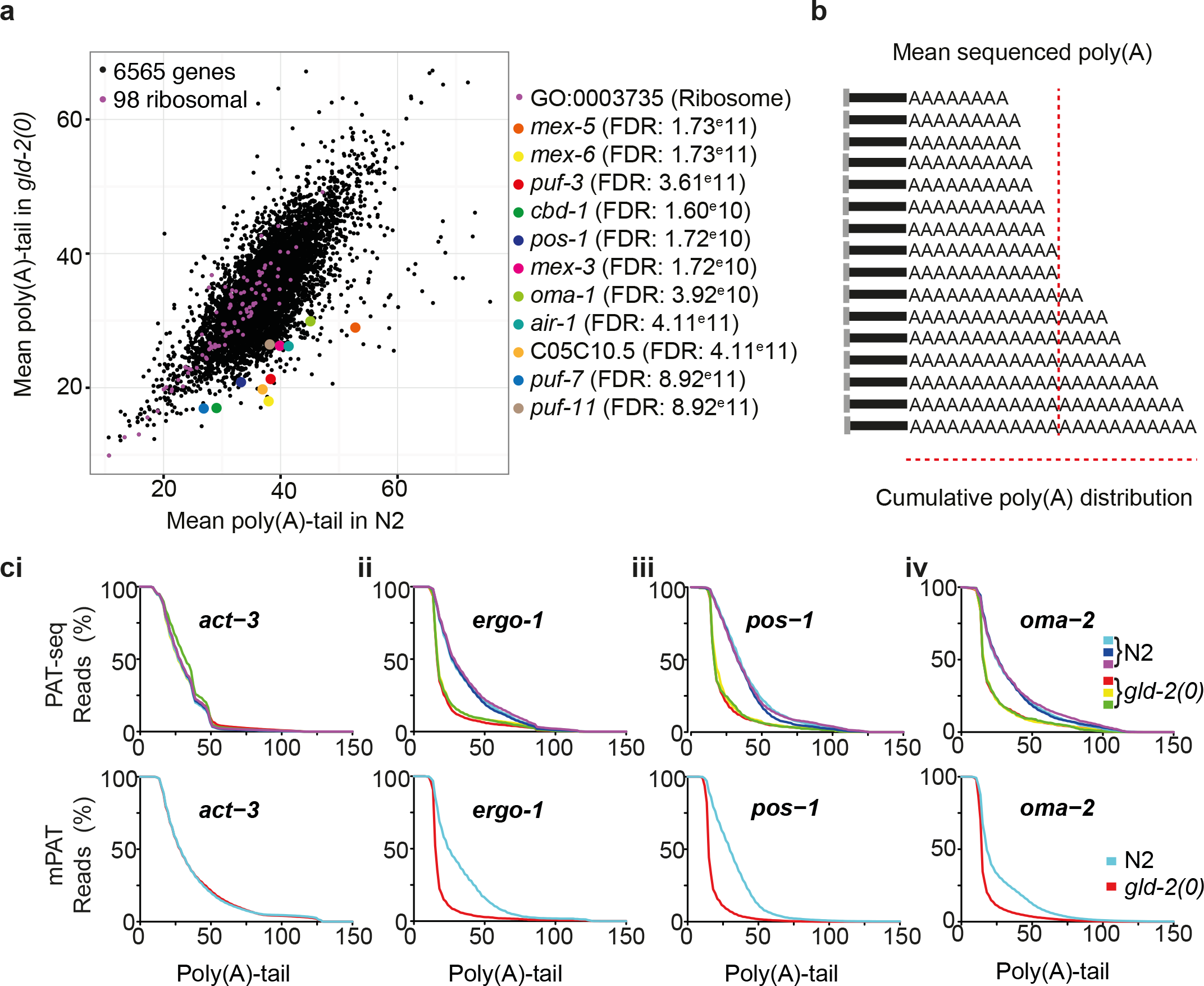
The measurement and visualization of polyadenylation-state change. a) To visualize individual GLD-2 targets an interactive tool was developed in the shiny framework. This can be used to highlight individual genes, groups of genes or GO terms. Genes indicated in purple (small dots) represent those associated with the GO term Ribosome (GO:0003735). The larger colored dots represent the 11 highest confidence GLD-2 targets, 9 of which are RNA-binding proteins. b) Schematic illustration poly(A)-tail length visualization. In comparison to simply reporting the average number of sequenced adenosines that terminate each read, visualization of the distribution of A-lengths measured can reveal biphasic and other skewed distribution patterns. c) Cumulative distribution of sequenced poly(A)-tails i) *act-3*; ii) *ergo-1*; iii) *pos-1*; and iv) *oma-2* as measured by PAT-seq (top panel) or as confirmed by mPAT (lower panel). The non-random cleavage pattern of RNaseT1 (after G residues) during PAT-seq NGS library prep can result in non-native distributions (eg *act-3*). These do not diminish the power to detect differences in adenylation-state between conditions, but the ‘shape’ of adenylation can be distorted. The corresponding mPAT assays (lower panels) support this conclusion.

The steady-state transcriptome represents adenylated mRNA reflective of a spectrum of metabolic states that include newly-transcribed (long-tails), silenced (short-tails), reactivated (long-tails), actively-translated (mid-range-tail) and ageing transcripts (shortening-tails). Because the steady-state length represents a sum of these activities, we reasoned that visualization of the length-distribution might be more informative than a simple numerical descriptor such as the mean or median sequenced length, which may hide complex distribution patterns (Fig 2b; (27). We therefore generated an interactive visualization tool to analyze the cumulative distribution of poly(A)-lengths for any gene of interest. This visualization approach highlights that the ‘shape’ of adenylation is highly reproducible and can be transcript-specific (Fig 2ci-iv). The *act-3* transcript, for example, displayed a highly uniform distribution of lengths that was independent of GLD-2, whereas the RNA-binding proteins *ergo-1*, *pos-1* and *oma-2* each showed a robust difference in length-distribution between the three wild-type and *gld-2(0)* transcriptomes analyzed.

To validate both adenylation-shape and distribution-change we applied a multiplexed high-throughput sequencing version of the ePAT assay, referred to as mPAT ((28) see methods). Twenty-nine gene-specific forward primers were used in a multiplexed and nested PCR reaction compatible with directional sequencing on the Illumina MiSeq platform. The sequenced poly(A)-distribution by this approach was highly similar to that recorded by PAT-seq (Fig 2ci-iv lower panel; and additional Fig 1a & b). With a multiplexed digital output for both read-count and sequenced A-residues, this is a significant technical advance to the traditional gel-based poly(A)-tests that depend on interpretation of changes in smear-distribution of PCR amplicons (eg. Fig 1d).

Since cytoplasmic adenylation is widespread among germline transcripts, we searched for any functional significance of gene ontology (GO) categories associated with GLD-2 target mRNA. As expected, enriched categories included those linked to germline biology, such as embryonic development, cell-cycle, cell-death, negative regulation of gene-expression and germ granules (known as P granules in *C. elegans*). The categories that were significantly under-represented included those related to signal-transduction, transcription factors, immune and neurological systems and genes encoding membrane proteins (additional Fig 2a and additional File 2).

Transcripts with the most dramatic poly(A)-shortening in our data encoded RNA-binding proteins (Fig 2a; additional Fig 1a & b). For example, MEX-1/3/5 & 6, OMA-1/2 and POS-1 are all RNA-binding proteins that are intimately involved in the 3’ UTR-mediated spatiotemporal control of post-transcriptional gene expression (13, 26, 29, 30). We therefore asked if auto-regulation through cytoplasmic polyadenylation was a general feature of transcripts encoding RNA-binding proteins. This was not the case however, as the trend for adenylation-state difference for transcripts annotated as RNA-binding in WormBase was similar to the global trend (additional Fig 2b). Because the GO searching identified germline function among the significantly short-tailed transcripts, we asked next if GLD-2 mediated extension of mRNA poly(A)-tails was a general feature of all mRNA within the germline, by overlay of the previously identified gene-lists for either the spermatogenic or oogenic germline (31) (additional Fig 2c). While there was a general trend toward slightly longer poly(A)-tails for spermatogenic, and slightly shorter for oogenic transcripts than the global PAT-seq average (see dashed lines), the data did not support the idea that germline mRNA are default targets of cytoplasmic polyadenylation. Finally, we searched for overlap between GLD-2 targets previously identified by physical RIP-chip of GLD-2 and RNP-8 (20). We expected enrichment of a defined sub-set of our GLD-2 targets specifically regulated by RNP-8. However, of the 365 genes shared between datasets, only a small proportion of genes were among those we identified as GLD-2 targets (additional Fig 2d). This observation can only be satisfactorily explained by the significant technical differences between the studies: co-purified of RNA associated with GLD-2/RNP-8 complexes (20), whereas our data directly surveyed the transcriptome for changes in poly(A) tail length as a surrogate of GLD-2 enzymatic activity.

### GLD-2 target mRNA preferentially depend on CCF-1 for deadenylation

To be detected as a GLD-2 target in our assay required mRNA to be short-tailed in the mutant compared to the wild-type transcriptome. Our assumption was that the targets of cytoplasmic adenylation would first be deadenylated as part of their storage and translational silencing (4, 16, 17, 22). We expected that depletion of one of the five deadenylases encoded in the *C. elegans* genome would reverse the short-tailed phenotype of target mRNA in the *gld-2(0)* mutant background. Each deadenylase was therefore systematically depleted by selective isolation of larval stage one (L1) *gld-2(0)* mutants and RNAi feeding (Fig 3a). Using the *ergo-1* transcript as an exemplar, we conducted ePAT assays of total RNA isolated from *gld-2(0)* mutants after each deadenylase knockdown and compared them to wild-type worms. Only knock-down of *ccf-1* restored the poly(A)-distribution in *gld-2(0)* mutant worms to the wild-type distribution (Fig 3b). To confirm that this was true of the majority of GLD-2 targets, full PAT-seq analysis was performed for each RNAi sample. This showed a strong global-trend toward longer poly(A)-tails after *ccf-1(RNAi)* knock-down in *gld-2(0)* worms (additional Fig 3a, blue dashed line), with 656 transcripts significantly longer in *ccf-1(RNAi)* compared to control (RNAi) using a conservative cut-off (FDR < 0.01) (additional Fig 3a, red dashed line). This trend was in broad agreement with traditional bulk adenylation-state analyses that previously suggested that CCF-1 is the major germline deadenylase (32). RNAi knockdown of *panl-2* also resulted in a global shift toward longer poly(A)-tails (additional Fig 3b), but neither this nor the other deadenlyases tested reached statistical significance for individual genes using our stringent tests for significance. Examples of the cumulative poly(A)-tail length-distributions are shown (additional Fig 3c) and the full dataset for all knockdowns is provided for interactive exploration (14).

**Fig 3:**
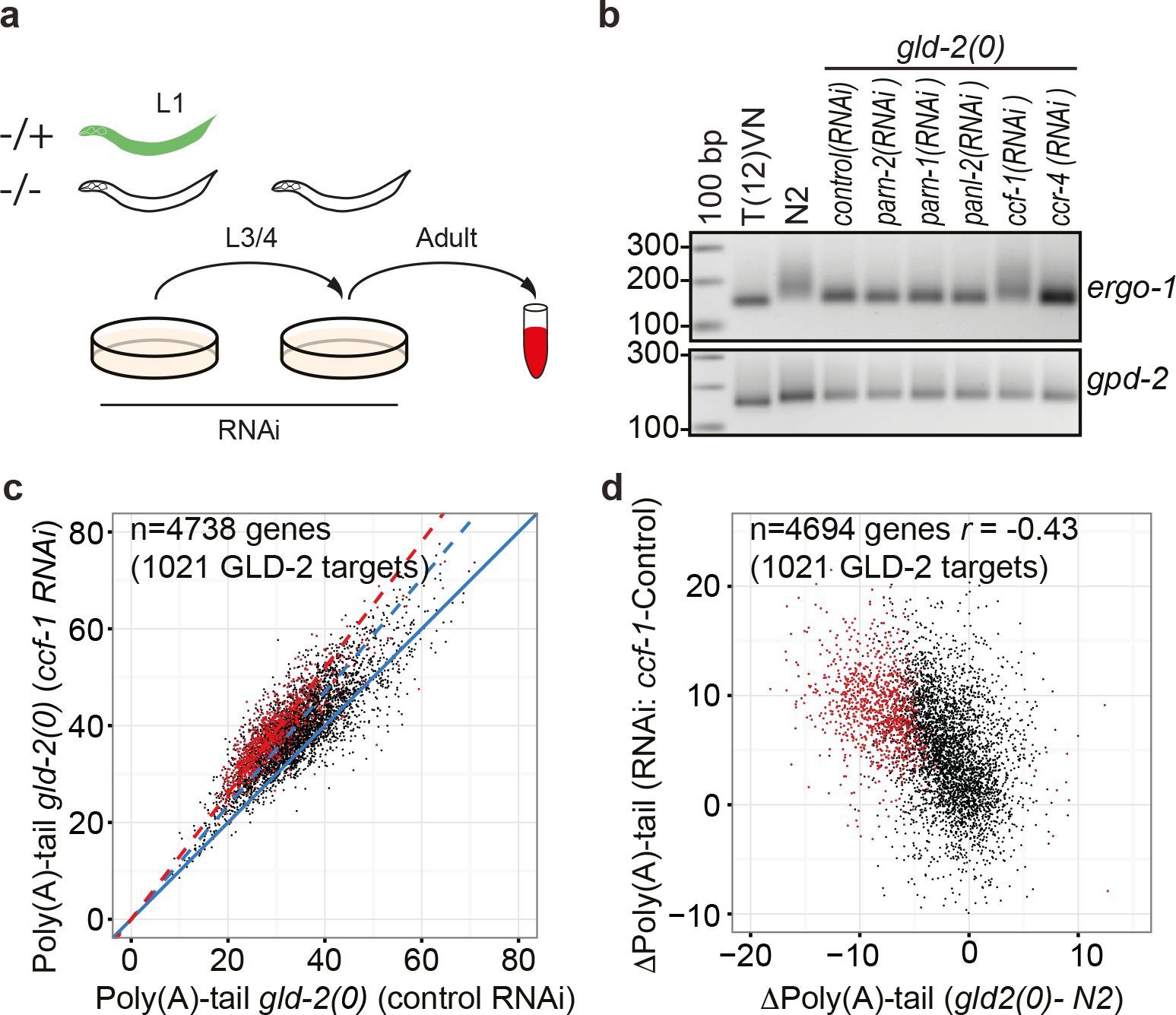
CCF-1 is the deadenylase responsible for poly(A)-tail trimming of GLD-2 targets. **a)** Schematic representation of the selection and treatment of *gld-2^−/−^* homozygous mutants for knockdown. Progeny from balanced heterozygous (*gld-2^−/+^*) worms were raised on RNAi plates. At the L3/4 stage of development, homozygous mutants (GFP-ve) were manually selected under fluorescent light onto fresh RNAi plates and harvested at the equivalent of 60 hours of development. **b)** Isolated RNA from each RNAi knock-down was analyzed by ePAT. The T(12)VN assay was prepared from a mix of the 6 RNAi samples. The tight migration of *ergo-1* ePAT amplicons in the control RNAi, indicates the shortening expected in the *gld-2(0)* mutant. Any recovery of the native (like N2 wild-type) smear of adenylation products indicates successful knock-down of the specific deadenylase required for shortening the tails of GLD-2 target transcripts. The *gpd-2* transcript is included as a control for unchanged adenylation-state. **c)** Comparison of the overall adenylation-state of the transcriptome of *ccf-1* (RNAi) versus control (RNAi) knock-down in *gld-2(0)* mutants. The significant GLD-2 target transcripts are indicated in red. Note that while there is a global increase in tail-length by *ccf-1(RNAi)*, (blue dashed line), the GLD-2 targets are more dramatically shifted (red-dashed line). **d)** There is a correlation (Pearson’s r=-0.43; *p*-value < 2.2e^−^16) between CCF-1 targets and GLD-2 targets. The difference between *gld-2 (0)* and wild-type adenylation-state and the loss of *ccf-1* (RNAi) versus control (RNAi) in *gld-2 (0)* mutants. The significant GLD-2 target transcripts are indicated in red. The accumulation of red in the top left-hand quadrant indicates that the transcripts most likely to be GLD-2 targets are also the most likely to depend on CCF-1 for shortening.

To confirm that GLD-2 target transcripts were specifically subject to targeted deadenylation by CCF-1, we compared their distribution pattern on plots showing the poly(A)-length changes between *ccf-1(RNAi)* and control RNAi (Fig 3c). The GLD-2 target transcripts displayed an overall longer adenylation-state (see red dashed line) than the transcriptome average (blue dashed line) after CCF-1 depletion. This was further supported by the inverse correlation observed between the adenylation difference between *gld-2(0)* and wild-type tail-length, and the difference between *ccf-1* and control RNAi strains (Fig 3d). The genes in the upper left-hand quadrant are those that were most shortened in *gld-2(0)* and are correspondingly the most lengthened by *ccf-1(*RNAi) in *gld-2(0)*, and were overwhelmingly enriched for our high-confidence GLD-2 targets (colored red). These data support a model where deadenylation is an early step in the metabolism of mRNA that are marked for cytoplasmic polyadenylation within the germline and/or later within the oocyte-to-embryo transition.

### 3’ UTR of GLD-2 target mRNA are longer, c-rich, and contain non-canonical poly(A)-motifs

What is it that marks a mRNA for silencing deadenylation and subsequent activation by re-adenylation? The PAT-seq approach identifies polyadenylation sites with exquisite sensitivity, which is confirmed by the high level of overlap with previously annotated positions of 3’ UTRs in *C. elegans* (33) and our wild-type transcriptome (additional Fig 4a). A common idea is that the length of 3’ UTR is correlated to the complexity of post-transcriptional regulation (34). Therefore, to ask if the 3’ UTRs of GLD-2 target transcripts differ in any global parameters compared to those that are apparently not targeted, we first asked if there was a relationship between 3’ UTR length and the change between wild-type and *gld-2(0)* adenylation state (additional Fig 4b). Interestingly, GLD-2 target mRNAs were associated with longer 3’ UTR, with an average of 307 versus 189 bases in GLD-2 target versus non-target 3’ UTR respectively (p≪0.0001). This finding was supported by trend inversion in *ccf-1(RNAi)* worms (additional Fig 4c).

We next asked if any linear regulatory sequence motifs could be identified in the 3’ UTR of GLD-2 target mRNAs. Motif searching revealed that GLD-2 target 3’ UTR were often associated with an alternative, non-canonical poly-adenylation sequences (most commonly AAUGAA), functions for which had not been previously identified. Testing for the frequency of each of the major reported adenylation site sequences (33), revealed that any of the alternative sites were more likely to be targeted by cytoplasmic polyadenylation than those with canonical AAUAAA sites (additional Fig 4d). The same sites were associated the transcripts having a *ccf-1(RNAi)* mediated rescue of adenylation in the *gld-2(0)* mutant background (additional Fig 4e). In an important caveat, a canonical recognition motif did not preclude an mRNA from being targeted for cytoplasmic polyadenylation. For example, the *gld-3* transcript is expressed as two isoforms with different 3’UTR (35). Both were targets of GLD-2 activity, but only the long isoform utilized the non-canonical poly(A)-signal (additional Fig 5). Together these data point toward a complex and fundamentally different mechanism of 3’-end metabolism for GLD-2 dependent transcripts.

**Fig 5:**
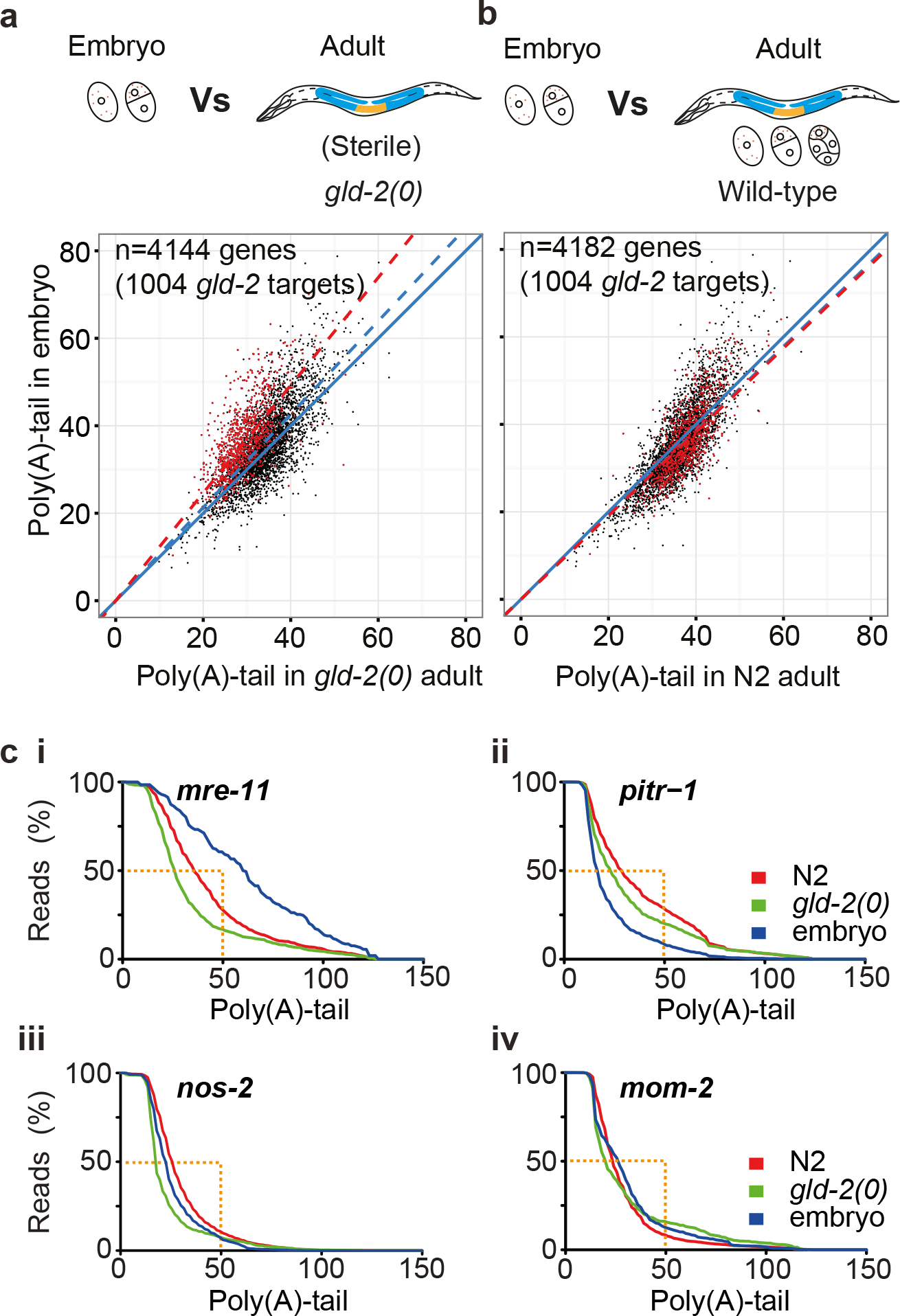
Polyadenylation-state changes reveals spatiotemporal control. **a)** Comparison between the poly(A)-tail distribution of the wild-type 1-2 cell embryonic transcriptome and the whole adult *gld-2*(0) mutant transcriptome, which, being sterile does not contain mature oocytes or viable embryos. The adenylation-state of transcripts present in both transcriptomes are indicated in the scatter plot. GLD-2 target transcripts are colored in red. **b)** Comparison between the poly(A)-tail distribution of the wild-type 1-2 cell embryo transcriptome and the wild-type whole adult worm transcriptome. Transcripts present in both transcriptomes are indicated, with the GLD-2 target transcripts in red. **c)** Individual GLD-2 target transcripts present with different adenylation states in the embryo (blue) versus the adult wild-type worm (red) or in *gld-2(0)* mutant (green). The dashed gold lines give a context to the absolute tail-lengths sequenced for each of the indicated transcripts. **i)** *mre-11* presents with the longest tail in the embryo, whereas **ii)** *pitr-1* harbors the shortest tail in the embryo, suggesting that its cytoplasmic adenylation is instead triggered earlier in germline development that decays in the embryo. The majority of the **iii)** *nos-2* and **iv)** *mom-2* transcripts are short-tailed in the conditions tested.

Foundational studies in vertebrate cytoplasmic polyadenylation suggested specific placement of motif for recruitment of cytoplasmic poly(A)-polymerase (11). Here, searching the high confidence GLD-2 targets for enriched motifs using Homer (36), returned only the AAUGAA motif described above as differential between the two groups. Reasoning that the RNA-binding proteins that recruit the GLD-2 enzymatic activity likely drive specificity, we next searched specifically for RBP motifs in target mRNA (Fig 4; additional Fig 6, and (14)). These revealed that while the GLD-2 target mRNA were enriched for the AAUGAA motif, and mRNA with an unchanged poly(A)-length distribution are enriched for the AAUAAA polyadenylation site motifs, the position of these sites relative to the site of cleavage does not differ (Fig 4c & d, additional Fig 6a). Surprisingly, a search for enriched mono, di and tri nucleotide sequences revealed that the 3’ UTRs of GLD-2 targets were cytosine-enriched (Fig 4e). The C-patch was most concentrated directly upstream of the poly(A)-signal (Fig 4e (lower panel); and additional Fig 6a).

**Fig 4:**
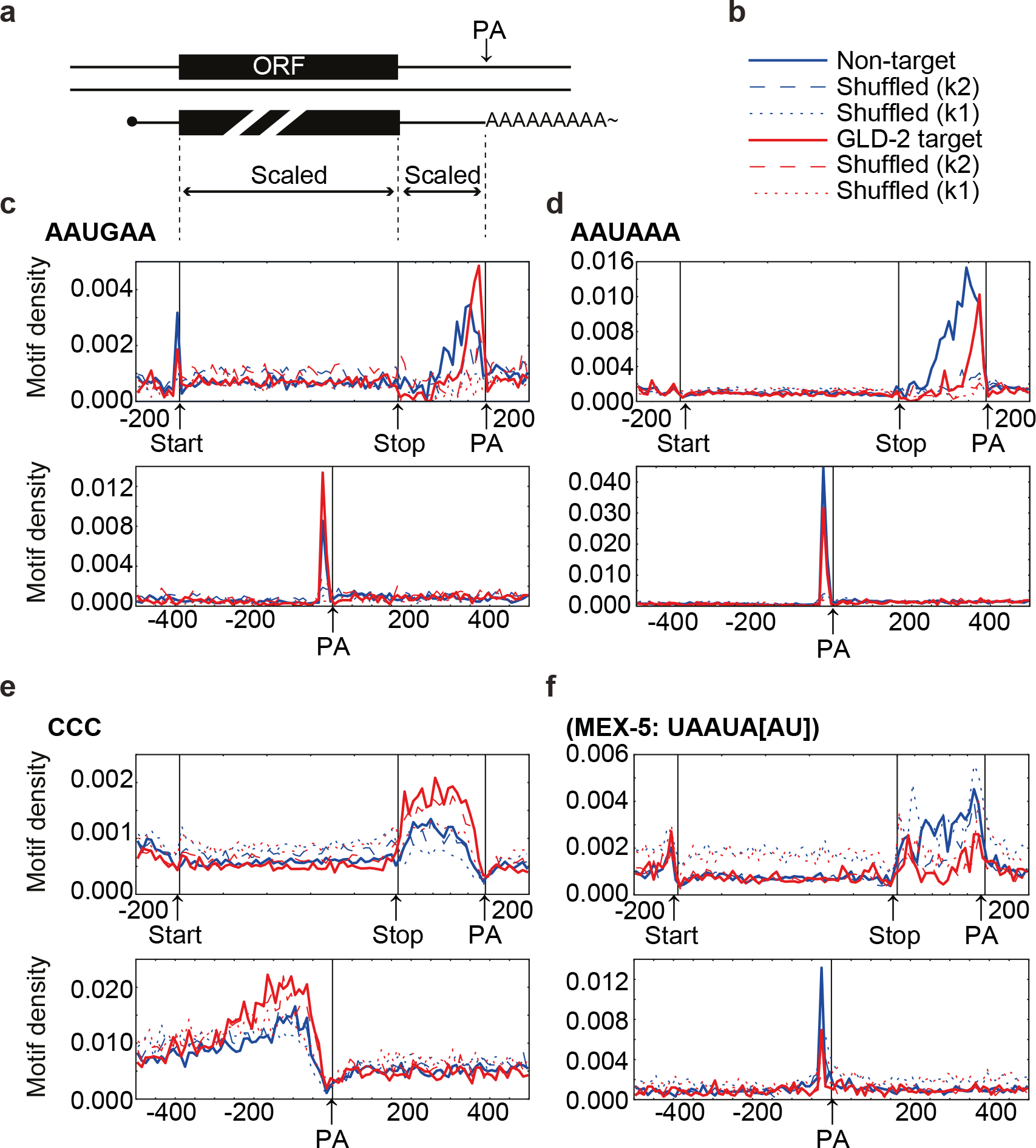
GLD-2 target 3’UTR are enriched for C-tracts and non-canonical Poly(A)-sites. **a)** Schematic to illustrate the scaling of information in the top panel of each of the figures below. Systematic search for motifs that differentiate the GLD-2 target set of transcripts from apparent non-targets can be found on the associated website (14) **b)** Graphs can be interpreted using the key where motif density in GLD-2 target transcripts are summarized by the red line and non-targeted transcripts are summarized by the blue line. The dashed lines of k1 and k2 show the result of searching for the motif on a locally shuffled version of the genome. When shuffling the genome, Individual bases were displaced with standard deviation of displacement by 10 bases. This preserves local nucleotide frequencies while scrambling any specific motifs. In the k2 version, this was done so as to also preserve dinucleotide frequencies. **c)** The non-canonical AAUGAA polyadenylation signal was enriched in GLD-2 target transcripts indicated by the increased height of the red peak just upstream of the site of polyadenylation (PA) (19.7% vs 12.8% p=9.59e^−7^). The apparent difference in distribution is an effect of the target-set having overall longer 3’ UTR. The motif is concentrated ~25 nucleotides downstream of the PA in both the target and non-target set when centered on the PA (lower panel). **d)** The canonical AAUAAA poly(A)-signal is enriched in non-target transcripts, indicated by the increased blue peak (58 % vs 42 %; p=3.38e^−17^) **e)** The 3’ UTR of the GLD-2 targets are C-rich. The 3’ UTRs unaffected by GLD-2 loss were 15.6% C, whereas the GLD-2 targets were 19.5% C. The CCC trinucleotide was strongly associated with 3’ UTR of GLD-2 targets (*p*=2e^−^65). **f)** CISBP-RNA gives UAAUAW (UAAUA[AU]) as the MEX-5 targeting motif. This element is positioned as an extension of the polyadenylation signal, i.e. UAAUAAA. This sequence was much more common in the non-target set (16.1%) than GLD-2 target 3’ UTR (8.35%) (p=4.62e^−12^).

A search for each of the motifs identified for twenty *C. elegans* RNA binding proteins in the CISBP-RNA database (37) confidently returned only the MEX-5 binding site (Fig 4f and (14)). However, identification of this motif appears to have been driven by a positional overlap with the canonical polyadenylation motif enriched in non-target 3’ UTR (Fig 4f, lower panel). Finally, we focused on the previously identified POS-1 motif (UA(U_2-3_)RD(N_1-3_)G) (38) which was commonly present in 3’ UTRs (50.5% in non-target, 71.9% in target). Being in the GLD-2 target set increased the odds of the POS-1 motif being present 2.5-fold, (*p*=5.0e^−29^). To check if this was merely due to the overall increased length of the GLD-2 3’ UTR, a logistic regression model predicting the presence of the POS-1 motif was constructed with a term for 3’ UTR length in addition to whether a gene was a GLD-2 target. The odds of having the POS-1 motif were still increased 1.52-fold by being a GLD-2 target in this model (p=6.5e^−6^) in GLD-2 targets. Therefore, the presence of the POS-1 motif in the GLD-2 group with a changed poly(A)-length distribution in *gld-2(0)* mutants was not simply due to increased 3’ UTR length (additional Fig 6b). It is important to note, however, that this is a highly redundant motif, and that POS-1 is itself expressed only asymmetrically in the early embryo (39, 40). Some GLD-2 targets having a legitimate POS-1 site might remain undetected in our data because they do not change in adenylation-state within the developmental stages analyzed or, only in subsets of under-represented cells.

### Staging polyadenylation-state control by comparison to the 1-2 cell embryonic transcriptome

Zygotic transcription only broadly commences at 4-cell stage in the worm embryo (2). Maternal RNA adenylation-state is thus likely to be dynamically regulated in the steps between its initial synthesis, the P-granules that deliver mRNA to the developing oocytes, and their activation during the oocyte-to-embryo transition. Therefore, we next profiled adenylation-state in 1-2 cell wild-type embryos (Fig 5). Comparison of the adenylation-state of the embryonic-transcriptome with the sterile *gld-2(0)* transcriptome, illustrates that the majority of shared mRNA that are short-tailed in the *gld-2*(0) mutant (red) are long-tail in the embryo and presumably activated for translation within the developing embryo (Fig 5a) supporting the GO classification of GLD-2 target transcripts (additional Fig 2a). On the other hand, comparison of the 1-2 cell embryo transcriptome with the wild-type transcriptome, showed that these same GLD-2 targets presented with a spectrum of lengths, some being longer in the total transcriptome and shorter in the embryos, and vice-versa. GLD-2 target transcripts that were shorter-tailed in the 1-2 cell embryo likely represent those that were activated earlier in development, that is, having been activated within the germline syncytium or oocytes and having undergone subsequent poly(A)-shortening with translation, aging and maternal RNA clearance in the embryo.

The differential adenylation-state between adult and embryo can reveal context specific regulation. Four GLD-2 targets that exemplify such control are *mre-11, pitr-1, nos-2* and *mom-2* (Fig 5 ci-iv). These transcripts each show a unique pattern of poly(A)-tail length distributions between the wild-type, *gld-2(0)* and embryo transcriptomes. The *mre-11* transcript encoding a conserved DNA recombination and repair protein is apparently expressed at its highest levels in first 30 minutes of embryonic development (41, 42) and this is supported by the broad distribution of poly(A)-tails in the embryo transcriptome (>50% having a sequenced length of > 50 A-residues). The *pitr-1* transcript, by contrast, is expressed mainly within the germline (43), and thus the poly(A)-distribution is shorter as it decays in the embryo, as compared to the two adult transcriptomes. The *nos-2* transcript is a GLD-2 target, but because it is active in both the stem cells of the germ line and in the later stages of the embryo, its poly(A)-tail does not reach adult lengths in isolated 1-2 cell embryos. Finally, *mom-2*, a member of the Wnt family of signaling proteins and required for gut specification is short-tailed in both the adult and the 1-2 cell embryo (see also additional Fig 1b). The spatiotemporal translation of *mom-2* was previously shown to depend on the combinatorial action of nine RNA-binding proteins (44) for expression in a single cell (P2) for specification of its sister cell (EMS) at the 4-cell stage of development (See Fig 1a). The length-distribution of *mom-2* in the wild-type adult and 1-2 cell embryo, and in the *gld-2(0)* adult, suggest that it is a target of highly restricted cytoplasmic adenylation. These data prompted us to ask; can spatially restricted transcripts be identified on the basis of their dependence on regulatory RNA-binding proteins for a wild-type adenylation profile?

### Asymmetric activation of maternal mRNA in the embryo

With a compendium of GLD-2 targets in hand, we reanalyzed previous PAT-seq data that examined the adenylation-state of the early embryo after RNAi knockdown of the GLD-2 anchor protein GLD-3 and selected 3’ UTR regulators ((13) and Fig 6a). In the previous report, asymmetric expression of a new regulator of endo-mesoderm development NEG-1 (repressor of endoderm fate in the AB cell) was shown to depend on GLD-3 for recruitment of GLD-2, and on POS-1 for repression in the P1 embryonic lineage. Focusing on early embryos depleted for either POS-1, MEX-5 or GLD-3 and comparing these to the matched wild-type embryos revealed that the majority of embryonic GLD-2 targets depended on GLD-3 to reach the wild-type polyadenylation profile (Fig 6b ii-xii). However, at the 2-cell stage, a binary switch is established that depends on MEX-5 and POS-1, two RNA-binding proteins that reciprocally regulate each other’s expression (13, 25). This repression is reflected in their dependence on each other to curtail cytoplasmic polyadenylation to the wild-type level (Fig 6b ii & iii), albeit *mex-5* also appears to depend on its own expression to reach the wild-type adenylation-state. In the absence of MEX-5, multiple mRNAs show extended poly(A)-tails (Fig 6b iv-vii) whereas other mRNAs are regulated by POS-1 (Fig 6b iii, viii-xi).

**Fig 6:**
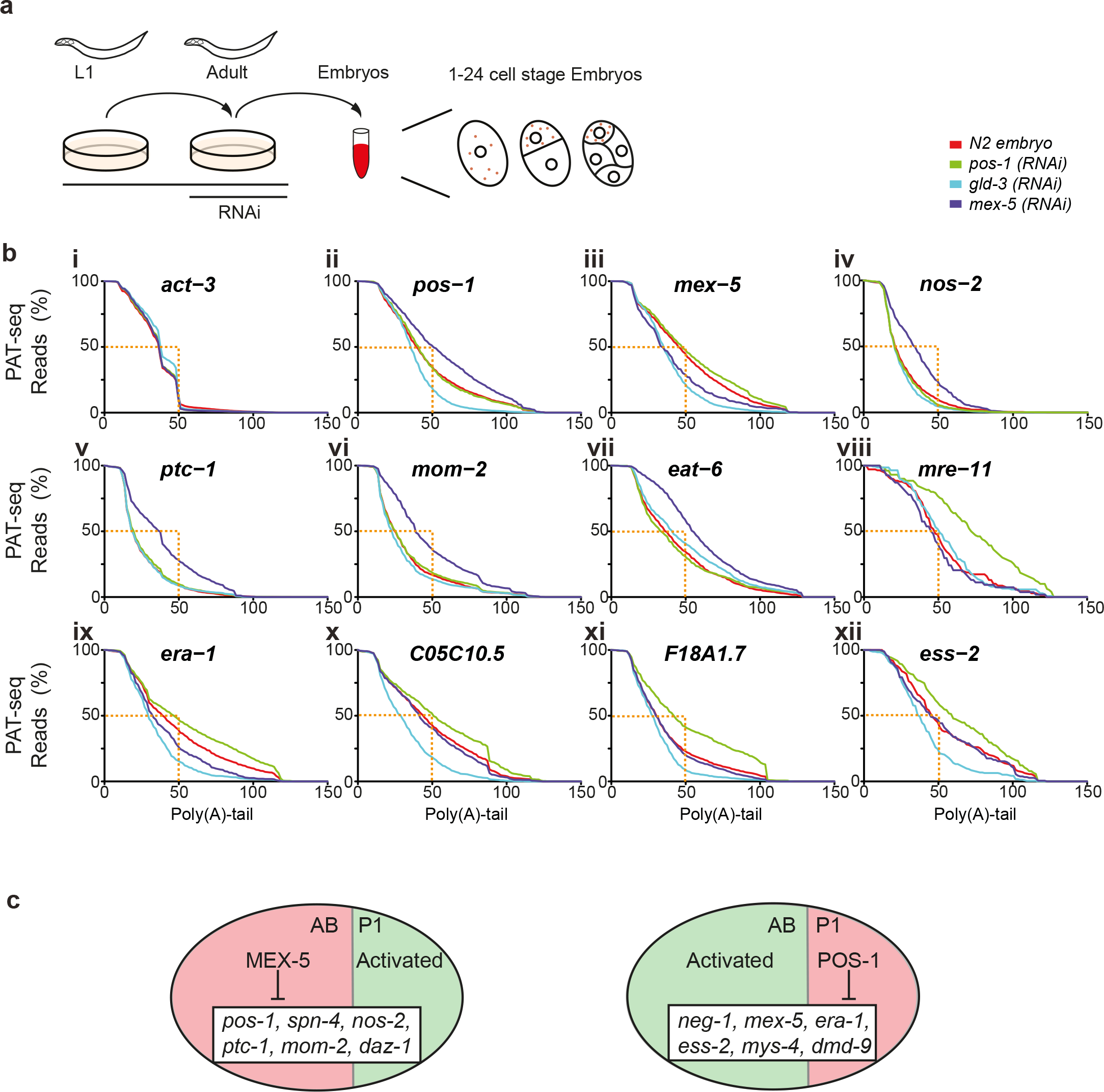
RNA binding proteins regulate the spatiotemporal control of GLD-2. **a)** Previously, PAT-seq data was collected from isolated 1-24-cell embryos after timed depletion of the RNA-binding proteins POS-1, MEX-5 or GLD-3 by RNAi (13). The data from the resulting embryo pools were reanalyzed here in the context of cytoplasmic polyadenylation. **b)** Cumulative distribution of the sequenced poly(A)-tails was plotted for the indicated transcripts. The lengths sequenced are given context by the dashed gold line. **i)** *act-3* is not a *GLD-2* target and is uninfluenced by the RNA binding proteins tested. The transcripts indicated in panels **ii-xii** are targets of GLD-2 mediated cytoplasmic polyadenylation, but differentially depend on GLD-3 (light-blue), the anchor for the GLD-2 mediated cytoplasmic polyadenylation in the oogenic germline, and the regulators of this activity POS-1 (green) and MEX-5 (purple) to maintain the wild type (red) adenylation-state. **c)** The adenylation-states observed can be understood when considered in the context of the 2-cell embryo where maternal transcripts are distributed between both the AB (presumptive ectoderm and neurons) and the P1 cell (presumptive endo-mesoderm and germline) but are differentially regulated. In the AB cell MEX-5 is expressed and represses the expression of the indicated transcripts. Whereas, POS-1 is expressed in the P1 cell and represses cytoplasmic polyadenylation of the indicated targets. The targets of POS-1 and MEX-5 are recognized by their hyperadenylation, due to aberrant cytoplasmic polyadenylation in both cells.

POS-1 and MEX-5 are major drivers of polarity in the first cell division by repressing cytoplasmic adenylation and thereby translation of target mRNA. However, these proteins are themselves asymmetrically localized, and therefore repress their targets cell-type specifically (Fig 6c). Depletion of either RNA-binding protein by RNAi resulted in cytoplasmic polyadenylation of maternal RNA in both cells of the 2-cell embryo stage of development. Searching the data for transcripts that were selectively hyper-adenylated after either MEX-5 or POS-1 depletion (similar to those shown in Fig 6b ii-xii), we identified many of the known mRNA encoding key factors required for correct embryonic patterning and multiple new candidate mRNAs that are likely poised for asymmetric expression in the early embryo (additional File 3). Importantly, while many of the 102 potential MEX-5 targets are known and include those critical for cell fate specification (eg *nos-1*, *ptc-1*, *mom-2*, *daz-1, apx-1* and *zif-1;* see also additional Fig 1a & b) less than half (22/52) of the transcripts that were repressed by POS-1 have been annotated with any function. These provide excellent candidates for specification of the AB-derived somatic lineages, the asymmetric programming of which is much less well understood than the germline lineages. Collectively these examples, and those that can be found by further interactive search of our data in the accompanying web-apps (14), are consistent with a model where MEX-5 controls AB fate and POS-1 the P1 fate by negative regulation of the cytoplasmic polyadenylation of target mRNA. However, we expect that the many additional 3’ UTR binding proteins will overlay this binary switch to further refine fate-decisions in the early embryo.

## Discussion

The development of technologies to detect changes to adenylation state between transcriptomes (12, 45, 46) has meant that the functional influence that poly(A)-tail length-control has on post-transcriptional gene expression networks is now open for detailed investigation. Here, we map RNA metabolism, through changes in adenylation-state of the transcriptome under multiple experimental conditions. By this approach we traced the selection, silencing and subsequent activation of maternal mRNA in space and in time. The driving question for this study was to understand if cytoplasmic polyadenylation is utilized in the germline as a regulatory switch for a few key drivers of early development or whether it represented a more global mechanism of regulatory action. Our data suggest that there are many targets of GLD-2 mediated poly(A)-extension. We identified >1000 high confidence targets via a change in adenylation state in adult *gld-2(0)* mutant worms (Fig 1). Yet, this number is likely an under-representation given that whole worms were profiled, and that some cell types or transient effects may have been under-represented. Although we identify a surprisingly large number of targets (~16% of the detected transcriptome), cytoplasmic polyadenylation is not likely to be a default activity unleashed on all germline transcripts. As indicated by the enriched gene ontologies (additional Fig 2), transcripts targeted by GLD-2 are not those required for general building projects, but instead encode the regulators and their targets that require precisely timed expression in time and space.

Cytoplasmic polyadenylation has been shown as critical for male fertility in the fruit fly (24) and by parallel evolution also in the mouse (47). A surprising finding in our work was the stabilization of short-tailed spermatogenic transcripts in the *gld-2(0)* adult suggesting a failure in normal RNA metabolism and clearance after the switch to an oogenic germline. This raises the possibility that GLD-2 is required for both activation of targets in the spermatogenic transcriptome (additional File 1) and possibly also of an as yet to be identified regulatory factor responsible for clearance of the spermatogenic mRNA. A failure of spermatogenic P-granules (48) to resolve their cargo prior to the switch to the oogenic germline, might explain aberrant transcript persistence and could contribute to the sterility of these animals (18). Candidates for regulatory factors required for clearance of the spermatogenic transcriptome are the CPEB proteins CPB-1 and FOG-1, with the transcript of at least the former being a clear GLD-2 target (14) and both proteins were shown by others to be required for the germline-switch (23). Alternatively, the persistence of spermatogenic transcripts might be explained by a failure to repress transcription of the male-specific transcriptome in the *gld-2(0)* mutant. Importantly, the adenylation-state data reported here provides a new resource to guide the study of the mechanisms that underpin the germline-switch.

The analysis of 3’ UTR sequences indicated that the 3’ UTR of GLD-2 target mRNA were overall longer, tended to utilize the non-standard hexa-nucleotide motifs, and had a different (C-enriched) sequence character (Fig 4; additional Fig 4-6). Thus, similar to the recent study comparing *wisp* mediated cytoplasmic polyadenylation and its impact on translation-state in *Drosophila* (49), no single motif appears to be the driver of cytoplasmic polyadenylation in the worm. This finding contrasts with the prevailing ideas surrounding CPEB1 binding to its cognate 3’UTR binding sites in vertebrates (50, 51), and suggesting that there is much still to be discovered regarding the control of cytoplasmic polyadenylation across evolution. However, consistent with our findings, a C-rich element was previously suggested as a recruitment element for cytoplasmic adenylation of the PP2Ac transcript in *Xenopus* oocytes (52). Moreover, a previous study of mRNA turnover during oocyte-to-embryo transition revealed that a polyC motif was both necessary and sufficient for maternal RNA clearance in *C. elegans* (53). Whether this represents a shared motif that switches function, triggering first timed cytoplasmic polyadenylation and then clearance of maternal RNA, or whether it simply represents a feature of this particular cohort of mRNA remains to be determined. Of the two polyC binding proteins proposed to mediate the decay of maternal mRNA, *hrpk-1* and *nova-1*, the former is itself a GLD-2 target, consistent with a role for GLD-2 in most aspects of temporally regulated germline RNA metabolism.

The advent of high throughput approaches such as PAT-seq, means that the details of how 3’ UTR elements converge to control mRNA sequestration, and subsequent timed reactivation is now open to interrogation. To be identified as a GLD-2 target in our study, the adenylation-state needed to change between *gld-2(0*) and wild-type worms. Nascent transcripts are first polyadenylated by the canonical nuclear poly(A)-polymerase (PAP-1) concomitant with 3’-end cleavage in the nucleus. Yet, we had previously observed that selected transcripts displayed a shortened poly(A)-tail in the *gld-2(0)* mutant (22, 54) suggesting that this initial polyadenylation was removed during P-granule storage of mRNA. The deadenylase PARN1 has been proposed to buffer GLD-2 activity in *Xenopus* oocytes (55). Here, however, only depletion of CCF-1 had an overt impact on the adenylation-state of GLD-2 targets. Both CCR-4 and CCF-1 are highly expressed in the germline and are present at all oocyte developmental stages, however, only CCF-1 induces sterility in *C. elegans* whereas CCR-4 knockdown worms are fertile, albeit with fewer offspring (56). The shared targets we observed by knockdown of CCF-1 in the *gld-2(0)* mutant (Fig 3) are consistent with a functional pairing of CCF-1 and GLD-2 in the linear narrative of transcript silencing and subsequent reactivation at the correct time and place in development.

The patterning of the early embryo begins with a cascade of auto-regulation by RNA-binding proteins. RNA-binding proteins with known (and yet to be discovered) roles in the patterning of the germline and embryo are among the most significantly short-tailed in the *gld-2(0)* mutant compared to adult worm transcriptome (Fig 2). Here, we show that the targets of such RNA-binding proteins can be identified and tracked by changes to adenylation-state after factor knock-down (Fig 6). Moreover, the sensitivity of the PAT-seq and mPAT approaches we have developed provide valuable new tools to distinguish subtle changes in polyadenylation. Here, we identified mRNA subject to differential adenylation after asymmetric cell division that is conveniently visualized by cumulative distribution plots. However, more subtle changes to adenylation-state can also be determined by such visualizations. For example, in a previous analysis in the fly embryo, as little as ~5% of a transcript population could be identified as differential between samples (27). We therefore propose that the experimental analysis tools are now in place to systematically mine RNA-driven cellular asymmetries in developmental and disease biology.

## Conclusions

Collectively our data suggest a model whereby GLD-2 regulates all stages of germline and early embryonic development. Starting with the precise switch from male to female germline, development and maturation of oocytes, and in the oocyte-to-zygote transition (Fig 7). The GLD-2 target mRNA whose expression is spatiotemporally regulated in the worm contain 3’ UTR encoded information that differentiates them early in their biogenesis from non-target mRNA. This information is encoded in longer 3’ UTR that utilize non-canonical polyadenylation signals and contain a C-rich 3’ UTR patches. We propose that this marks them for capture by P-granules abutting the nuclear pore complexes. These granules are then shed into the shared cytoplasm and within this context become deadenylated by CCF-1. GLD-2 activity is then selectively recruited by an exquisitely timed combination of anchoring proteins such as GLD-3 and regulatory proteins such as MEX-5 and POS-1. Importantly, we propose that this complex circuitry will be accessible by combinatorial depletion of RNA binding proteins and measure of polyadenylation-state.

**Fig 7:**
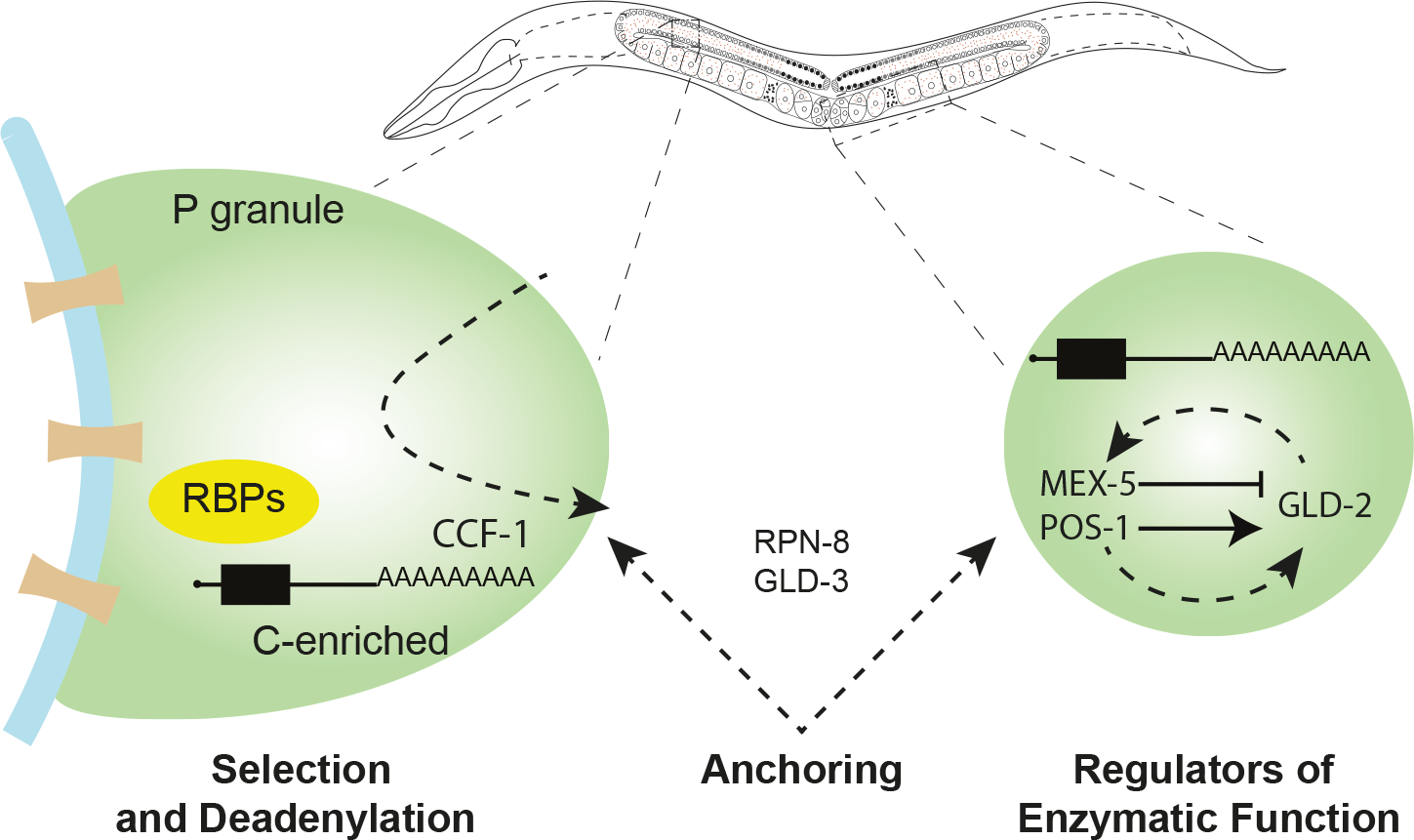
GLD-2 regulates the *C. elegans* germline and oocyte-to embryo translation by the sequential action of RNA-binding proteins that regulate its activity in space and time. Schematic model for the capture, transit and activation of GLD-2 target mRNA. Germline mRNA destined for GLD-2 mediated translational regulation are captured and deadenylated in the P-granules that abut the nuclear membrane. As these are shed from the Nuclear periphery of either masculinized or feminized germline, specific RNA binding proteins (RNPs) are required to bind target 3’UTR as an anchor for the GLD-2 enzymatic function (eg GLD-3, RPN-8). A second set of temporally active RBPs (eg POS-1 and MEX-1) regulate the enzymatic function of GLD-2.

## Methods

### *C. elegans* culture, growth and analysis

Strains used were Bristol strain N2 as wild-type, PRB106 (*gld-2(q497)/* hT2 [bli-4(e937) let-?(q782) qIs48]) and LD1101 (*cbp-3(tm1746)*) and were maintained using standard methods (57). The *gld-2(q497)* worms were maintained over the hT2 balancer chromosome that encodes a GFP marker. Homozygous *gld-2(q497)* worms are identified by the lack of GFP marker expression. RNAi was performed by bacterial feeding using *E. coli* HT115(DE3) strain carrying RNAi constructs targeting *parn-1, panl-2, ccr-4, parn-2*, and *ccf-1*. As an RNAi control (*pCB19)*, a fragment of an *Arabidopsis thaliana* gene that shares no homology with any *C. elegans* gene was used (58). For RNAi knockdown experiments, synchronized first stage larvae (L1) from *gld-2(q497)/ hT2 [bli-4(e937) let-?(q782) qIs48]*adults were placed on RNAi plates and at the L3 stage homozygous *gld-2(q497)* worms (non-GFP expressing) were picked to a new RNAi plate. Young adults (~60 hours) were collected and snap frozen in liquid nitrogen in biological replicate. For analysis of the early embryonic transcriptome, gravid adult N2 worms where dissected and 1-2 cell stage embryos collected (~100 per biological replicate) by aspiration and snap frozen in liquid nitrogen.

Dissection and immunostaining of *C. elegans* germlines was conducted as described in (59). Antibodies to CGH-1 and IFET-1 were used at 1/200 dilutions and slides were analyzed using an inverted Olympus 1 x 81/1 x 2-UCB microscope attached to an X-cite series 120Q fluorescent light box with a DAPI filter at 461nm. Images were obtained with a 40X oil immersion objective, using Cell-M software (Olympus, Tokyo, Japan).

### RNA analyses

To isolate total RNA ~100 worms per sample were collected in the lid of a 1.5 ml tube and snap frozen in liquid nitrogen. The frozen bead of worms was flash centrifuged and a volume equivalent to ~250 mg Zirconia beads (BioSpec Products) and 1 ml of Trizol (Invitrogen) was added. The samples were pulsed for 30 seconds at top speed in a mini-beadbeater-8 (BioSpec Products). The handling from this step forward was according to the Trizol product specifications, with 200 μl of chloroform being added directly to the tubes, containing the zirconia beads. Once the clear aqueous phase was transferred to a fresh tube, 2 μl of co-precipitant glyco-blue (Ambion) was added to ensure efficient RNA precipitation. The ePAT and TVN-PAT assays were performed using 1 μg of total RNA exactly as previously described (22). All relevant primers including those for mPAT are listed in Additional File 4.

### High Throughput Sequencing (HTS)

PAT-seq analysis was performed as previously described (12). For each library, 1 μg of total RNA was used as input. For the mPAT (<u>m</u>ultiplexed <u>P</u>oly(<u>A)</u>-<u>T</u>est) assays, cDNA synthesis was exactly as previously described for ePAT (22), except that the anchor primer used to tag the 3’ ends was Illumina compatible, and a 23 base extension was appended 5’-end of each gene-specific primer (28). Nested PCR reactions introduced further (P5/P7) Illumina compatible elements. In the first step, each individual primer was suspended to 100 μM, then stoichiometrically pooled (i.e total stock oligo mix remaining at 100 μM) for a multiplexed first round PCR reaction using the mPAT anchor primer as a reverse primer and the pooled mix of 29 forward primers (Additional File 1). Conditions used for amplification were as for ePAT except that in the first PCR reaction, the volume was 100 μl, with 25 μl of diluted (1/5) cDNA as template. Five cycles of amplification were used. The first round PCR amplicons were recovered by a column clean-up step using NucleoSpin^®^ PCR Clean-up columns (Macherey-Nagel). To promote removal of oligonucleotide primers, the binding buffer was diluted and used at ½ strength as suggested in the product specifications. First-round amplicons were eluted from the columns in 49 μl of dH_2_O pre-warmed to 60 °C. For the second round of PCR, all of the first-round amplicons were re-amplified with commercial ScriptSeq (Epicentre) indexed reverse primers (i.e. one index per sample) and the PAT-seq universal sequencing primer. A proportion (10 %) of each library was visualized by 2% high-resolution agarose gel electrophoresis to ensure equal library amplification. The individual libraries were then pooled and enriched by column cleanup as above. Pooled libraries were sequenced using a MiSeq V2 reagent kit and 300 cycles according to the manufacturer’s specifications at the Monash University sequencing facility, Micromon. Note: Primers were designed (where possible) within ~100 bases of the polyadenylation site to ensure unique amplification of 3’-end while maintaining sufficient sequencing cycles to determine polyadenylation. Note: A lab-ready protocol for mPAT is included as Additional File 5

### Data analysis

PAT-seq and mPAT NGS data were analyzed using the tail-tools bioinformatic pipeline (12). The tail-tools software package (12, 60) was used to align reads to the ce10 reference genome. This software calls gene expression and poly(A)-sites. The average poly(A)-tail length was calculated from reads terminating in non-templated A-tracts at these sites. For each gene, the site with the most reads in wild-type was taken as the primary poly(A)-site. Such sites were allowed to be up to 1000 bases downstream of the annotated end of the gene. This allowed a 3’ UTR region of each gene to be accurately defined, and the sequence of these regions to be examined. The GEO accession for the sequencing data is GSE68002.

Our data can be analyzed by a series of dedicated interactive tools (14). The tools included are two interactive web apps for differential tests of DGE and poly(A)-tail length-distribution implemented here as part of an R-Shiny online interface (61). For the Poly(A) cumulative distribution visualization tools, adenylated reads associated with each gene were extracted from the bam files using the Rsamtools R package (62). Users can select a gene of interest in an optional set of samples, and the application extracts all reads with a poly(A)-tail and plots the distribution of lengths sequenced. All data showing the cumulative distribution of poly(A)-tails in this manuscript were generated with this app and modified (font, size etc.) in Adobe Illustrator. The poly(A)-tail and gene expression plotter, displays search genes of interest, gene lists or GO categories in the context of a scatter-plot comparing the *gld-2* and wild-type transcriptomes (as in Fig 2a). The user chooses which condition (gene expression or tail-distribution) to plot against each other. For user-guided visualization of scatter plots the count files are filtered according to the user’s input. The expression counts were normalized to CPM using the varistran package (63). Inputs that enable the user to interactively highlight specific genes of interest or to highlight genes associated with a particular GO Term utilize the biomaRt package (64). Any resulting figures and tables selected can be downloaded. Alternatively, all PAT-seq datasets can be interactively searched using the ‘Degust’ tool for RNA-seq exploration analysis and visualization (65) within the dedicated web companion to this research. Note: the significant GLD-2 and CCF-1 targets were based on this output.

### 3’ UTR analysis for enriched Motifs

To mine 3’ UTR for motifs, genes with a significantly shorter poly(A)-tail in the *gld-2(0)* than in wild-type were identified using the Fitnoise software package (60). This adapts the limma (66) approach of using linear models with moderated variance to the problem of detecting differential tail length. For this analysis, genes were required to have at least one *gld-2(0)* mutant sample and one wild-type sample having at least 50 reads containing a poly(A)-tail. Thus 994 target genes (FDR 0.01 cut-off) were compared to a neutral set (1953 genes), in which the average tail length differed by at most 2 A-residues between the *gld-2(0)* mutant and wild-type. In both of these data sets, genes were required to have a CDS of at least 200 bases and a 3’ UTR of at least 20 bases. The average length of the 3’ UTR in the neutral set was 188.8, and in the differential set the average was 307.2.

We searched for motifs using Position Probability Matrices (PPM) downloaded from the CISBP-RNA database (37). A sequence was counted as a match if its probability of being randomly generated using the PPM was within 0.25 times the probability of the most likely sequence to be generated from the PPM. We also searched for specific motifs identified in the literature: POS-1 motif: UAU(2,3)[AGU][ACGU](1,3)G, canonical PAS: AAUAAA, alternative PAS: AAUGAA, reverse complement canonical PAS: UUUAUU, MEX-5: UUUUUU, UUUUUUUU, UAAUA, UAAUA[UA], UAAUAAA The density of cysteines in the 3’ UTR was enriched in the (994 gene) GLD-2 target set as compared to the (1953 gene) non-target set. Note that the cut-off here was slightly more stringent than that used by the differential tests where 1065 statistically significant GLD-2 targets were reported. For further details please see the interactive search tool for ‘motifs’ in the accompanying web app (14).

## Declarations

### Ethics approval and consent to participate

Not Applicable

### Consent for Publications

Not applicable

### Availability of data and material

PAT-seq NGS data is available in raw formats (GSE68002 this study) and (GSE57993 from Elewa *et al*., 2015) as well as by interactive web-based search tools available on our web companion to this manuscript .

### Authors’ contributions

PRB and THB designed the experiments, interpreted the results and wrote the manuscript. PFH, AAB, ADP, KT, EH and DRP contributed bioinformatics analyses. AAB, GR, AS, SM and GMD contributed experimental data.

## Acknowledgements

We thank members of the Beilharz Lab for constructive comment on the manuscript and the Monash Technology Platforms for technical support in next generation sequencing and bioinformatics. Melissa J. Curtis is acknowledged for technical support with the mPAT assay.

## Additional Figures

**Additional Figure 1a & b: Validation of PAT-seq identified changes to adenylation state.**

Differential poly(A)-tail length-distributions identified by PAT-seq were validated for 23 transcripts by mPAT approach to multiplexed ePAT amplicon resequencing on the Illumina MiSeq Platform.

**Additional Figure 2: Cytoplasmic polyadenylation is not a default feature of germline transcripts.**

**a)** The Gene Ontologies enriched and depleted in GLD-2 targets. The search was done by inputting the 1692 (FDR < 0.05) into the FuncAssociate GO term finder (67). For further visualization of how such Gene Ontology groups distribute in our data see the poly(A)-tail and gene expression plotter app that accompanies this manuscript (14).

**b)** Scatter plot based on the difference in mean sequenced poly(A)-length between the *gld-2(0)* and wild-type transcriptome (as in Fig 1d). Transcripts that are annotated as ‘RNA-binding’ in Wormbase are indicated (red). These do not distribute in an obviously different manner to the transcriptome on a whole (see red vs blue dashed lines).

**c)** The transcripts previously identified in the isolated oogenic or spermatogenic germline (31) are colored in red and green respectively. The dashed lines indicate the average distribution of each subset and genes are colored accordingly.

**d)** The transcripts identified as RPN-8/GLD-2 interacting by RNA ImmunoPrecipitation (RIP) (20) are colored red. Of the 361 RPN-8/GLD-2 RIP-transcripts overlapping with our data, there was only a marginal difference in the mean tail-length change over the global difference of the transcriptome (see red vs blue dashed lines).

**Additional Figure 3: CCF-1 is the deadenylase responsible for shortening tails of GLD-2 targets.**

**a)** Plotting the mean tail-length between the *gld-2(0)*; *ccf-1(RNAi)* and *gld-2(0)*; control (RNAi) transcriptomes shows lengthening of a large proportion of the transcriptome (blue-dashed line). The proportion of transcripts that reach statistical significance for tail-length change from the replicate experiment are indicated in red. The dashed red line indicates the trajectory of the significant subset.

**b)** Plotting the mean tail-length between the *gld-2(0); panl-2(RNAi)* and *gld-2(0)*; control (RNAi) shows lengthening of a proportion of the transcriptome (blue-dashed line) as expected. However, these did not reach statistical significance by the stringent statistics implemented in our analysis pipeline.

**c)** Examples of cumulative tail-length distributions in the control *gld-2(0)*; *ccf-1(RNAi)*, *gld-2(0)*; control (RNAi) and 1-2 cell embryos. For some transcripts, the embryonic transcriptome presents with a poly(A)-distribution identical to the *gld-2(0);* control (RNAi) eg. *act-3, daz-1, gpd-4*. For others that are activated for translation in the embryo, the tail is longer (*ergo-1, pos-1, mex-5, puf-3, oma-1*). The *gpd-4* transcript is embryonically expressed, and is among the statistically significant GLD-2 targets we identify. Its distribution here is longest in the *gld-2(0)*; *ccf-1*(RNAi), however, since the tail is the same length in the 1-2 cell embryos as in *gld-2(0)*; control (RNAi), its cytoplasmic polyadenylation might only occur at later time point in development (see also *mom-2* Fig 4C). These data can be explored further in the associated web-apps.

**Additional Figure 4: The 3’ UTR of GLD-2 targets are longer and utilize divergent PAS.**

**a)** PAT-seq identifies the 3’-UTR poly(A)-junction transcriptome-wide at nucleotide resolution. Comparison of the genomic location of transcript adenylation-sites between the wild-type transcriptome measured here and in previous work (33) differ only in instances where there are discrepancies between the way 3’-ends were called. In our pipeline, the major adenylation site (i.e. the one with most reads) was chosen, whereas Mangone, Manoharan (33) chose the site most distal to the stop codon.

**b)** GLD-2 targets tend to have longer 3’ UTR. Plotting the difference in adenylation state between the *gld-2(0)* and wild-type transcriptome against the length of the 3’ UTR, indicates a broad distribution, but with a distinct correlation between longer 3’ UTR and larger adenylation-state difference. Curves were fitted using the LOESS method as implemented by the R function loess, with quadratic fitting and a span of 75%. The length difference is between the averages is 118.4 bases, 95% CI [104.0, 132.7], p≪0.0001. The shaded area indicates the 95% confidence interval. Significantly shorter tails (FDR ≤ 0.01) and poly(A)-tails with a difference of at least +−2 As.

**c)** As in **b**, except that recovery of tail-length after *ccf-1*(RNAi) is compared to the control RNAi treatment in *gld-2(0)* mutant worms. The longer the 3’ UTR the bigger the difference CCF-1 effect on tail-length.

**d)** The density of the canonical and non-canonical polyadenylation elements in the transcriptome as a function of tail-length difference between *gld-2(0)* and wild-type worms (error bars show 95% confidence interval of the mean). The canonical AAUAAA signal is not precluded from being a target (see also Fig 4d), but the non-canonical sites tend to have a bigger difference in adenylation-state change in the conditions determined here.

**e)** As in **d**, except recovery of tail-length after *ccf-1(RNAi)* was compared to the control RNAi treatment in the *gld-2(0)* mutant strain for enrichment of polyadenylation signals.

**Additional Figure 5: The canonical PAS does not preclude cytoplasmic polyadenylation.**

**a)** All eukaryotic transcriptomes show abundant evidence for alternative polyadenylation (APA). We found surprisingly little evidence for either differential expression of 3’UTR isoform or poly(A)-tail length-change between alternative 3’ UTRs. The *gld-3* transcript is an example of an mRNA that undergoes APA to generate alternate protein isoforms. In the IGV screen-shot, PAT-seq and mPAT reads are shown aligned to the transcriptome to highlight the position of aligned reads relative to the annotated 3’ UTR.

**b)** The two *gld-3* isoforms use different poly(A)-signals as indicated yet both are subject to GLD-2 mediated cytoplasmic polyadenylation. Albeit the shape of adenylation is slightly different, with the distribution poly(A)-tails of the distal isoform tending to be slightly longer than those associated with the proximal isoform. This distribution difference is reflected both in the original PAT-seq data and in the mPAT validation data (lower panel). Such difference in the shape of adenylation might become useful once combined with knock-down of RNA-binding proteins to understand the different function of the isoforms in time and space.

**Additional Figure 6: The character of GLD-2 target 3’ UTR.**

**a)** Analysis of the nucleotides surrounding the PA of GLD-2 target vs non-target transcripts. K-mer (k=1) analysis of the sequence 300bases up and 100 bases downstream of the PAS in the total transcriptome (top) shows the distribution of all bases. The next two panels show the difference in base composition between the target and non-target 3’ UTR. The target RNA tended to be T-rich with a cluster of C (C-patch indicated by *) just before the poly(A)-signal. The non-target 3’ UTR tend to be G rich with the region before the poly(A)-signal being more likely to be T-rich. Searching for 6-mers reveals an enrichment of poly(A)-signals, with AATAAA being the most abundant. Searching for differences, the AATGAA non-canonical sequence again being enriched in the GLD-2 target set.

**b)** The POS-1 motif is abundant in both target and apparently non-target 3’ UTR (see increased motif density in the 3’ UTR region). However, given that only the GLD-2 targets that changed in adenylation-state within the transcriptome of the adult were detected in our data, any transcripts that were targeted for cytoplasmic polyadenylation in an under-represented cell-population may not have been detected with a changed poly(A)-tail in our dataset.

## Additional Tables

**Additional File 1:** A list of the retained spermatogenic transcripts in the adult *gld-2(0)* worm. **Additional File 2:** A companion file to Additional Fig 2a. The file contains the full list of Gld-2 targets (FDR < 0.05), the wormbase IDs, and GO enrichment that were identified using FuncAssociate 3.0. tool.

**Additional File 3:** A list of the predicted targets of POS-1 and MEX-5 mediated repression of adenylation based on deregulated poly(A)-tail length after RNAi in early embryo.

**Additional File 4:** The list of oligonucleotide primers used in the study

**Additional File 5:** A lab-ready protocol for the mPAT method for multiplexed poly(A)-tail length discrimination.

